# Impairment of sugar transport in the vascular system acts on nitrogen remobilisation and nitrogen use efficiency in Arabidopsis

**DOI:** 10.1101/2022.09.14.507887

**Authors:** Beate Hoffmann, Emilie Aubry, Anne Marmagne, Sylvie Dinant, Fabien Chardon, Rozenn Le Hir

## Abstract

The carbon (C) and nitrogen (N) metabolisms have long been known to be coupled, and this is required for adjusting nitrogen use efficiency. Despite this intricate relationship, it is still unclear how a deregulation of sugar transport impacts N allocation. Here we investigated, in Arabidopsis, the consequences of the simultaneous downregulation of the genes coding for the sugar transporters *SWEET11*, *SWEET12*, *SWEET16,* and *SWEET17* genes on various anatomical and physiological traits ranging from the stem’s vascular system development, plant biomass production, seed yield, and N remobilisation and use efficiency. Our results show that intracellular sugar exchanges mediated by SWEET16 and SWEET17 proteins specifically impact the vascular development but do not play a significant role in the distribution of N. Most importantly, we showed that the double mutant *swt11swt12*, which is also impacted in the vascular development, displays an improved nitrogen use efficiency and nitrogen remobilisation to the seeds. In addition, a significant negative correlation between sugar and amino acids contents and the inflorescence stem radial growth exists, highlighting the complex interaction between the maintenance of C/N homeostasis and the inflorescence stem development. Our results thus deepen the link between sugar transport, C/N allocation and vascular system development.

**Highlight:** The disruption of genes coding for SWEET11 and SWEET12 sugar transporters negatively impacts the stem development but improves the plant nitrogen use efficiency and nitrogen remobilisation to the seeds.

## Introduction

Carbon (C) and nitrogen (N) are essential and limiting elements for plant, animal, and microorganism growth. In plants, the tricarboxylic acid (TCA) cycle is the primary source of C skeletons required for ammonium assimilation and is linked to the amino acid metabolism by the glutamate dehydrogenase, thus bridging C and N metabolisms (Huppe and Turpin, 1994; Hodges, 2002). In addition, N plays a significant role in C metabolism due to its function in Rubisco synthesis. At the same time, C compounds are essential for N absorption, nitrate reduction, N2 fixation, and amino acid metabolism to generate C skeletons, metabolic energy, and reductants (Baslam *et al.*, 2021). To successfully reproduce, plants must constantly adjust their C and N contents. Such modulation can be achieved at various levels of control, ranging from the modulation of nutrient assimilation proteins’ activity and the control of nutrient transport to a variety of mechanisms controlling the expression of genes encoding the proteins involved in nutrient metabolism, transport, and signaling. This has been exemplified in many studies which focused on improving C/N metabolism and transport to reach higher plant yield (Braun *et al.*, 2014; The *et al.*, 2021; Marmagne *et al.*, 2022).

To reach the seeds, sugars and amino acids, the primary products of C and N metabolisms, are transported through the vasculature (i.e., xylem and phloem) (van Bel, 2021). It is now well established that the main C pools are delivered long distance from the phloem, whereas the N pools are provided from both xylem and phloem. A third significant transport pathway is the lateral transfer of sugars and amino acids between phloem and xylem and *vice versa* (Tegeder and Masclaux-Daubresse, 2018; Aubry *et al.*, 2019). To ensure an appropriate distribution of sugars and amino acids at the whole plant level, nutrients can move through the plasmodesmata (symplasmic pathway), even if experimental proof of such a transport pathway for amino acids is still lacking (Kim *et al.*, 2021a), or by specialised transporters (e.g. SUGAR WILL EVENTUALLY BE EXPORTED TRANSPORTERS (SWEET), USUALLY MULTIPLE ACIDS MOVE IN AND OUT TRANSPORTERS (UmamiTs), SUCROSE TRANSPORTERS (SUC), and the AMINO ACID PERMEASES (AAPs) (Ladwig *et al.*, 2012; Müller *et al.*, 2015; Tegeder and Masclaux-Daubresse, 2018; Xue *et al.*, 2022). Overall, the delivery of nutrients from cell to cell and long-distance encompasses an intricate network between the metabolism products and their transport pathways that *in fine* creates source (e.g. leaves and stems) and sink organs (e.g. seeds, roots). The functional characterisation of these specialised transporters proteins shows that they are expressed in different plant organs (roots, leaves, stem, seeds), tissues (xylem, phloem, parenchyma cells, mesophyll cells) and subcellular compartment (plasma membrane or tonoplast), suggesting that tight coordination of their transport activities is needed for every step of plant development and growth. In this respect, genetic modifications have been used to understand the contributory role of sugar and amino acid transporter genes in increasing the C and N nutrition of seeds in crop and non-crop species (Wingenter *et al.*, 2010; Zhang *et al.*, 2010; Lu *et al.*, 2020; Grant *et al.*, 2021). For instance, the concurrent overexpression of amino acid permease AAP1 and sucrose transporter SUT1 in pea increased the nutrient fluxes from source to sink, resulting in increased seed number and protein content (Grant *et al.*, 2021). Nonetheless, despite the need to develop new strategies to improve plant yield and quality, experimental data regarding the impact of deregulation of sugar transport on N allocation or amino acids transport on C allocation is still scarce (Perchlik and Tegeder, 2018; Lu *et al.*, 2020).

In this work, we proposed to fill this gap by evaluating consequences of a deregulation of members of the SWEET sugar transporter family on biomass production, seed yield and N allocation in Arabidopsis. Based on our previous work, we focused on the concurrent downregulation of *SWEET11*, *SWEET12*, *SWEET16* and *SWEET17* genes shown to be expressed in the inflorescence stem (Le Hir *et al.*, 2015; Aubry *et al.*, 2022). Indeed, this organ constitutes an obligatory step for nutrients to be transported from leaves to seeds and is therefore of interest when studying whole plant C and N allocation. Overall, the analysis of the *swt11swt12, swt16swt17* double mutants and the *swt11swt12swt16swt17* quadruple mutant allowed us to establish a link between the sugar transport between the different vascular cell types, nitrogen use efficiency (NUE) and nitrogen remobilisation to the seeds (NRE).

## Materials and Methods

### Plant material and growth conditions

In order to obtain the quadruple mutant *sweet11-1sweet12-1sweet16-4sweet17* (hereafter referred as *swt-q*), we crossed the *sweet11-1sweet12-1* (hereafter referred as *swt11swt12*) double mutant (Le Hir *et al.*, 2015) with the *sweet16-4sweet17-1* (hereafter referred as *swt16swt17*) double mutant (Aubry *et al.*, 2022). Homozygous plants were genotyped using gene-specific primers in combination with a specific primer for the left border of the T-DNA insertion (Supplementary Table S1). By using RT-PCR analysis, we confirmed that full length transcripts of *SWEET11*, *SWEET12*, *SWEET16* and *SWEET17* cannot be amplified in the *swt-q* mutant line (Supplementary Fig. S1 and Supplementary Table S1 for primers sequences). To synchronize germination, seeds were stratified at 4°C for 48 hours and sown in soil in a growth chamber in long day conditions (16 hours day/8 hours night and 150 μE m^−2^ s^−1^) at 22/18°C (day/night temperature) with 35% relative humidity. Plants were watered with PlantProd nutrient solution twice a week (Fertil, https://www.fertil.fr/).

### GUS staining

The lines expressing pSWEET11:SWEET11-GUS or pSWEET12:SWEET12-GUS (Chen *et al.*, 2012) and pSWEET16:SWEET16-GUS or pSWEET17:SWEET17-GUS (Guo *et al.*, 2014) in Col-0 background were used to assess SWEET11, SWEET12, SWEET16 and SWEET17 expression pattern on seven-week-old plants grown in the greenhouse. The histochemical GUS staining was performed according to Sorin et al. (2005). Inflorescence stems subjected to GUS staining were then embedded in 8% (w/v) agarose and sectioned with a Leica VT100S vibratome (Leica, https://www.leica-microsystems.com/). Sections were counterstained for lignin by phloroglucinol staining (Pradhan Mitra and Loqué, 2014). Pictures were taken using a Leitz Diaplan microscope equipped with an AxioCam MRc camera and the ZEN (blue edition) software package (Zeiss, https://www.zeiss.com/).

### Inflorescence stem growth and sample preparation

The main inflorescence stem height was measured with a ruler at 45 days after sowing (DAS) and the stem diameter was measured either on Image J with the Feret diameter tool (see next paragraph) or with a digital caliper at the bottom of the stem. At 45 DAS, 1 to 2 cm segment was taken at the bottom part of the stem. Stem segments were embedded in 8% agarose solution and sectioned with a VT100 S vibratome (Leica, https://www.leica-microsystems.com/). Cross-sections were stained with a FASGA staining solution prepared as described in Tolivia and Tolivia (1987) for morphometric analysis of the different stem tissues.

### Morphometric analysis of the stem tissues

Stained inflorescence stem cross-sections were imaged under an Axio Zoom V16 microscope equipped with a Plan-Neofluar Z 2.3/0.57 FWD 10.6 objective (Zeiss, https://www.zeiss.fr/microscopie/). For each section, several parameters were measured: stem diameter, stem area, vascular system area (combining the area of all the vascular bundles), pith area, the phloem area (combining the phloem area of all vascular bundles), the xylem area (combining the xylem area of all vascular bundles) and the interfascicular thickness. The area of the other tissues (including epidermis, cortex and interfascicular fibres) was then calculated by subtracting the sum of the pith, phloem and xylem areas to the stem area. All parameters were measured using the Image J software package (https://imagej.nih.gov/ij/). For the same sections all the vascular bundles (VBs) were photographed individually using a confocal laser scanning microscope and morphological analysis of the xylem were performed as described in Le Hir et al. (2015). For each VB, the morphological segmentation allowed to access to the number of xylem cells (xylary fibres and xylem vessels) as well as their cross-sectional areas. Cells with a cross-sectional area comprised between 5 to 150 μm^2^ were considered to be xylary fibres and cells with a cross-sectional area greater than 150 μm^2^ were considered to be xylem vessels. The sum of all xylem cell cross-sectional areas was then calculated to give the total xylem cross-sectional area. The average xylary fiber and xylem vessel area was calculated by dividing the total xylem cross-sectional area by the number of each cell type.

### Quantification of soluble sugars, starch and total amino acids

Main inflorescence stems of wild type, *swt11swt12*, *swt16swt17* and *swt-q* mutants, without lateral stems, siliques and flowers, were harvested in the middle of the day (8 hours after the beginning in the light period), frozen in liquid nitrogen and ground with a mortar and a pestle. Soluble sugars and starch were extracted from 50 mg of powder from an individual stem and quantified by enzymatic method as described in Sellami et al. (2019). Total amino acids quantification was performed according to the protocol of Rosen (1957) and using Glutamine as a standard. Nine biological replicates coming from two independent experiments were analysed.

### Nitrogen and carbon percentage measurement and ^15^N labeling experiment

The different genotypes were grown in a growth chamber as described above. Around 30 DAS, while plants are still at the vegetative stage, 1 ml of a 10 mM nitrate solution containing 10% of ^15^NO_3_ was applied on the soil close to the plants collar. Then, plants were let to grow until the end of their cycle and harvested once all seeds were matured and the rosette dried. Plants were separated in different samples: (i) rosette, (ii) stem (including the main and lateral stems, cauline leaves and empty dry siliques), and seeds (total seeds). As plants were grown in soil, we could not harvest the roots without losing a large part of them. The dry weight of the different samples was determined. Then samples were ground to an homogenous fine powder and a subsample of 1000 to 2000 μg was precisely weighted in tin capsules to determine the C and N percentages (C% and N%) and the ^15^N abundance using an elemental analyzer (FLASH 2000 Organic Elemental Analyzer, Thermo Fisher Scientific, Courtaboeuf, France) coupled to an isotope ratio mass spectrometer (delta V isotope ratio mass spectrometer, Thermo Fisher Scientific, Courtaboeuf, France) calibrated using international reference (caffeine, IAEA-600, Vienna, Austria). The ^15^N abundance along with the following parameters: Harvest Index (HI), N allocation in rosette, N allocation in stem, N allocation in seeds (NHI), Nitrogen Use Efficiency (NUE = NHI/HI), ^15^N allocation in rosette, ^15^N allocation in stem, ^15^N allocation in seeds (^15^NHI) and Nitrogen Remobilisation Efficiency (NRE) were calculated as described previously (Jasinski *et al.*, 2021). The NRE index is calculated as the ratio between the ^15^NHI (percentage of ^15^N in seeds) over the plant HI (NRE = ^15^NHI/HI). Finally, the proportion of N coming from remobilisation or post-flowering uptake was calculated as described in Marmagne et al. (2022). The absolute quantity of C contained in each sample was defined as QtyC = DW x C%. The following formulas were then used to evaluate the parameters related to the C fluxes:

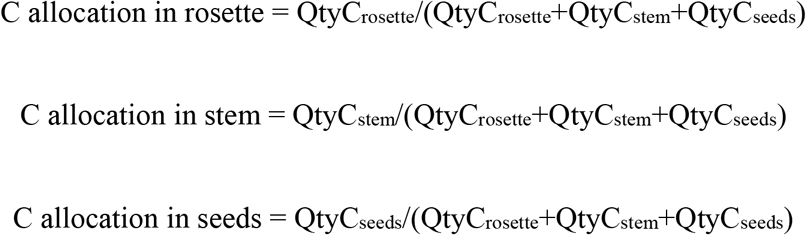

### RNA isolation, cDNA synthesis and RT-PCR

RNAs were prepared from the main inflorescence stem from 7-week-old plants grown as described above. Samples were frozen in liquid nitrogen before being ground with a mortar and a pestle. Powders were stored at −80°C until use. Total RNA was extracted from frozen tissue using TRIzol® reagent (Thermo Fisher Scientific, 15595-026, https://www.thermofisher.com) and treated with DNase I, RNase-free (Thermo Fisher Scientific, EN0521, https://www.thermofisher.com). cDNA was synthetized by reverse transcribing 1 μg of total RNA using RevertAid H minus reverse transcriptase (Thermo Fisher Scientific, EP0452, https://www.thermofisher.com) with 1 μl of oligo(dT)18 primer (100 pmoles) according to the manufacturer’s instructions. The reaction was stopped by incubation at 70 °C for 10 min. Full-length PCR was performed by using primers spanning the complete CDS for each gene tested (Supplementary Table 1).

### Statistical analysis

One-way ANOVA combined with a Tukey’s comparison post-test was done using R (version 4.0.2) and Rstudio (version 1.4.1103) software. A *P*-value <0.05 was considered as significant. Spearman correlations were realized using R with adjusted p-values calculated with the Holm’s method. Principal component analysis was performed using the ‘FactoMineR’ package of R (Le *et al.*, 2008). The least-square means were calculated using the R package ‘emmeans’.

## Results

It is now established that modifications of the facilitated transport of sugar at the plasma membrane or the tonoplast lead to defects in the Arabidopsis shoot and root development (for review see Xue *et al.*, 2022). However, to what extent perturbations of sugar exchanges at both the plasma membrane and the tonoplast lead to defects in the plant growth and development still needs to be explored. To address this question, we produced and characterised the *swt11swt12swt16swt17* quadruple mutant (*swt-q*) along the corresponding *swt11swt12*, *swt16swt17* double mutant lines (Le Hir *et al.*, 2015; Aubry *et al.*, 2022).

### The inflorescence stem growth and development are affected in quadruple sweet mutant line

Alike *swt11swt12*, the inflorescence stem of *swt-q* was shorter and thinner (by about 30%) than wild-type stem (Fig. 1A-C). Moreover, as previously described the stem of the *swt16swt17* was thinner but not shorter than that of the wild type (Fig. 1A-C) (Aubry *et al.*, 2022). The stem height to stem diameter ratio of both double mutants and the *swt-q* mutant was similar to that of the wild-type plants (Fig. 1D).

**Fig. 1.**
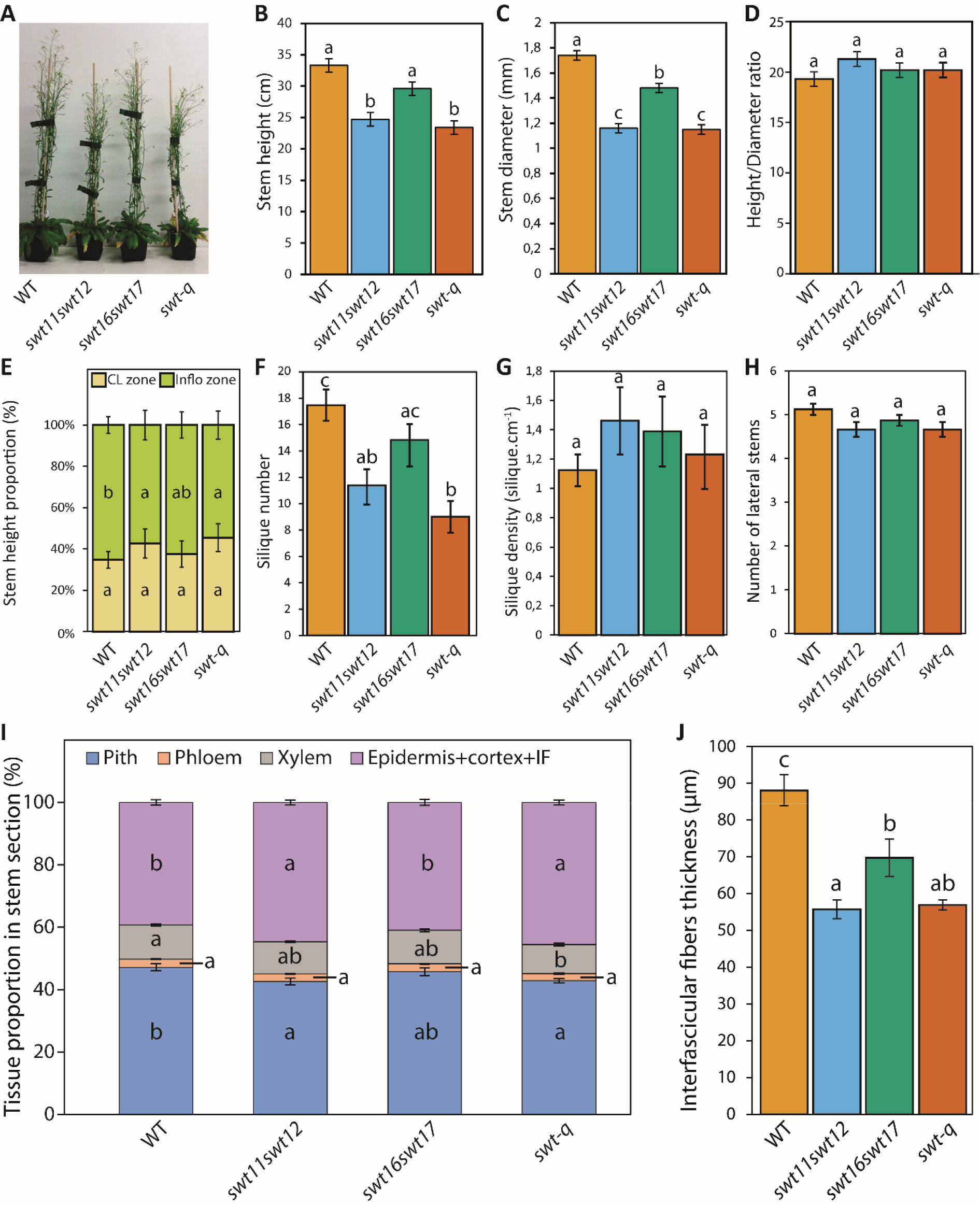
Main inflorescence growth and development is altered in the quadruple *sweet* mutant line. Photographs of a representative plant of wild-type, *swt11swt12*, *swt16swt17* and quadruple mutants taken at 45 DAS (A). Barplots showing the main inflorescence stem height (B), diameter (C) and the stem height-to-stem diameter ratio (D). Least-square means from three independent cultures ± SE (n=17 for each genotype). Letters indicate statistical difference between genotypes according to an ANOVA and Tukey’s comparison post-test. Proportion of the stem occupied by the “cauline leaf zone” (CL zone) and the “true inflorescence zone” (Inflo zone) (E). Barplots showing the silique number (F), the silique density (G) and the number of lateral inflorescence stems (H). Stacked barplot showing the distribution of tissues (epidermis+cortex+interfascicular fibres (IF), pith, phloem, and xylem) within an inflorescence stem cross-section (I) and barplot showing the interfascicular fibers thickness (J) among the different genotypes. Means ± SE (n ≥ 8 plant for each genotype). Letters indicate statistical difference between genotypes according to an ANOVA and Tukey’s comparison post-test.

We further characterised the inflorescence stem growth and measured after 45 days after sowing the portions of the stem corresponding to the cauline leaf zone, and to the “true inflorescence” (Pouteau and Albertini, 2009), the number of siliques, the density of siliques, and the number of lateral inflorescences (Fig. 1E-H). We observed that, in wild-type plants, the zone which carries the cauline leaves and the lateral inflorescence stems represents about 35% of the total inflorescence stem height while about 65% is occupied by the true inflorescence zone (Fig. 1E). Interestingly we observed that the proportion of this latter is reduced in the *swt11swt12* and *swt-q* mutants compared to wild-type plants (Fig. 1E). This phenotype is accompanied by a reduced number of siliques in the same genotypes (Fig. 1F). Nonetheless, the silique density was not significantly different between all genotypes studied (Fig. 1G). Finally, we did not observe any differences in the number of lateral inflorescence stems in any mutant lines compared to wild-type plants (Fig. 1H). We also explored the tissue distribution within the stem cross-section by measuring the area occupied by the pith, phloem, xylem and outer tissues (i.e. epidermis, cortex and interfascicular fibres) as well as the thickness of the interfascicular fibres (Fig. 1I-J). In our growth conditions, the pith represents about 47% of the wild-type stem cross-sectional area while 14 % is occupied by the vascular system (phloem and xylem). Finally, 39% of the total stem area is occupied by the epidermis, cortex and interfascicular fibres. The results are in accordance with those previously published by Paul-Victor and Rowe (2011). Interestingly the tissue organization changed in the different mutant lines (Fig. 1I). Indeed, a significant decrease of the proportion of xylem tissue was measured in the *swt-q* mutant compared to the wild type while no change in the proportion of phloem tissue was measured in any of the lines (Fig. 1I). Moreover, a decreased proportion of pith tissue along with an increased proportion of the outer tissue layers was measured in the *swt11swt12* and *swt-q* mutants. Additionally, the thickness of the interfascicular fibres was significantly smaller in all the mutant lines compared to the wild type (Fig. 1J). Therefore, the stem radial growth phenotype could be associated to a defect in the stem tissue distribution with less xylem, pith and interfascicular fibres.

Altogether these results show that both *swt11swt12* and quadruple mutants display similar phenotype, suggesting that adding perturbations of the cytosol-vacuole sugar exchanges (*via* disruption of the *SWEET16* and *SWEET17* expression) does not further affect the main inflorescence stem growth and development.

### Interplay between stem growth parameters and nutrient content

Both *swt11swt12* and *swt16swt17* double mutants have been shown to accumulate sugars in rosette leaves and inflorescence stem (Chen *et al.*, 2012; Gebauer *et al.*, 2017; Aubry *et al.*, 2022). Moreover, it has been demonstrated that the content of metabolites such as sucrose, glutamine and starch is negatively correlated with rosette biomass production (Meyer *et al.*, 2007; Sulpice *et al.*, 2009). Here, we tested the correlation between stem growth parameters and stem nutrient content in the different genotypes by measuring the main inflorescence stem diameter and height along with the total soluble sugars (sucrose, glucose and fructose), starch, and total amino acids content on the same samples (Figs. 1 and 2). A significant increase of soluble sugars, starch and amino acids contents was measured in the *swt11swt12* and *swt-q* mutant lines compared to wild-type (Fig. 2). Additionally, we show that *swt16swt17* stems tends to accumulate more soluble sugars compared to wild-type stems (Fig. 2). We then computed correlation coefficients between these parameters (Table 1). Interestingly, a negative correlation was shown between the inflorescence stem diameter/height and the total amino acids content as well as between the stem diameter and the soluble sugars and starch contents (Table 1). Additionally, a positive correlation between inflorescence stem height and diameter as well as a positive correlation between soluble sugars and starch contents were calculated (Table 1). These results suggest that the SWEET-dependent stem radial growth phenotype is associated with sugar and amino acid contents in this organ.

**Table 1.** Correlation matrix showing interrelationship between stem growth parameters and nutrients content. Spearman correlations with associated adjusted p-values calculated with the Holm’s method (n= 40 observations).

| **Variables** | Stem height | Stem diameter | Amino acids content | Soluble sugars content | Starch content |
| --- | --- | --- | --- | --- | --- |
| Stem height |  | 0.69 | -0.78 | -0.29 | 0.03 |
| Stem diameter | p<0.0001 |  | -0.69 | -0.51 | -0.45 |
| Amino acids content | p<0.0001 | p<0.0001 |  | 0.49 | 0.13 |
| Soluble sugars content | p=0.2033 | p=0.0058 | p=0.0071 |  | 0.49 |
| Starch content | p=0.8356 | p=0.0155 | p=0.8158 | p=0.0071 |  |

**Fig. 2.**
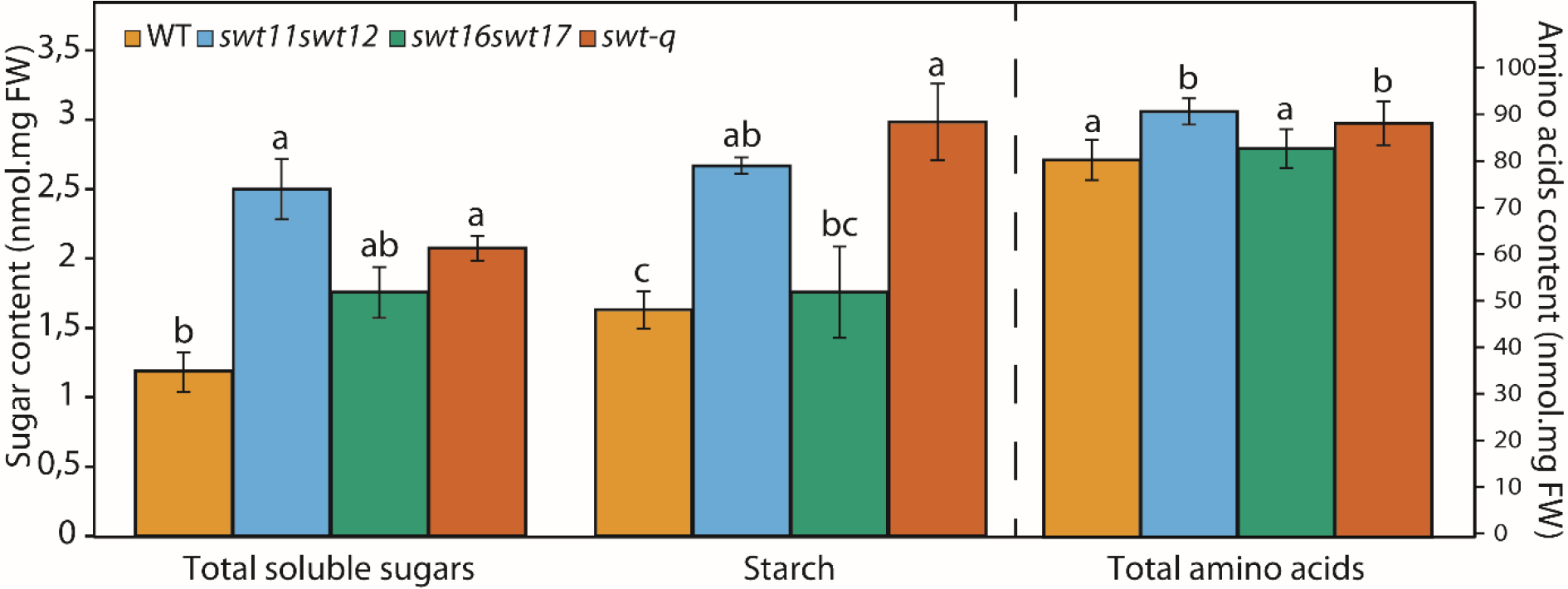
Soluble sugars, starch and amino acids accumulate in the main inflorescence stem of multiple *sweet* mutants. Barplots the total soluble sugars, starch and total amino acids content of wild type, *swt11swt12*, *swt16swt17* and quadruple mutants. Least-square means from two independent experiments +/− SE are shown (n=9 for each genotype). Different letters indicate significant difference according to a one-way ANOVA (Tukey’s test, p<0.05).

### Plant biomass production and seed yield are impaired in sweet mutant lines

Up to now, effects of down or up-regulation of *SWEET11*, *SWEET12*, *SWEET16* and/or *SWEET17* have been observed at an organ-specific level in rosette, stem, root or seeds (Chen *et al.*, 2012, 2015; Chardon *et al.*, 2013; Klemens *et al.*, 2013; Guo *et al.*, 2014; Le Hir *et al.*, 2015; Valifard *et al.*, 2021; Aubry *et al.*, 2022). Here, we wanted to obtain a more comprehensive view of the consequences of mutation in these genes on the aboveground organs at the end of the plant development. For that purpose, the biomass produced by rosette leaves, stem (including the main, lateral, secondary stems and the siliques envelopes) and seeds were assessed along with the analysis of C and N contents in each organ (Fig. 3). Consistently with previous report (Chen *et al.*, 2012), we show that rosette dry weight (DW) is significantly reduced in the *swt11swt12* double mutant compared to the wild type by about 45% (Fig. 3A). The same decrease in rosette biomass is observed for the *swt-q* mutant while no significant effect of mutation in *SWEET16* and *SWEET17* is measured (Fig. 3A). Interestingly a significant gain of biomass (about 40%) is measured only in the stem of the *swt16swt17* double mutant (Fig. 3A). Consistently with fewer siliques (Fig. 1F), the seed yield was reduced in both *swt11swt12* and *swt-q* mutant compared to wild-type plants (Fig. 3A). We also estimated the C and N allocated to each organ in the 4 genotypes. In wild-type plants, N was allocated at 6%, 37%, and 57% to rosette, stem, and seeds respectively (Fig. 3B). In *swt11swt12* and *swt-q* mutants, the N distribution was modified in all organs tested, with a significant decrease of N allocation in rosette and seeds while a significant increase of N allocation in the stem in both genotypes (Fig. 3B). C was allocated at 3.5%, 33% and 63.5% to rosette, stem and seeds respectively in wild-type plants (Fig. 3C). A significant decrease in C distribution was measured in the seeds of the *swt11swt12* and *swt-q* mutants while a significant increase of C was allocated to their stem (Fig. 3C). Finally, the *swt16swt17* double mutant displayed an intermediate phenotype for N and C allocation between WT and the other mutants (*swt11swt12* and *swt-q*) but no statistical difference was measured compared to the wild type.

**Fig. 3.**
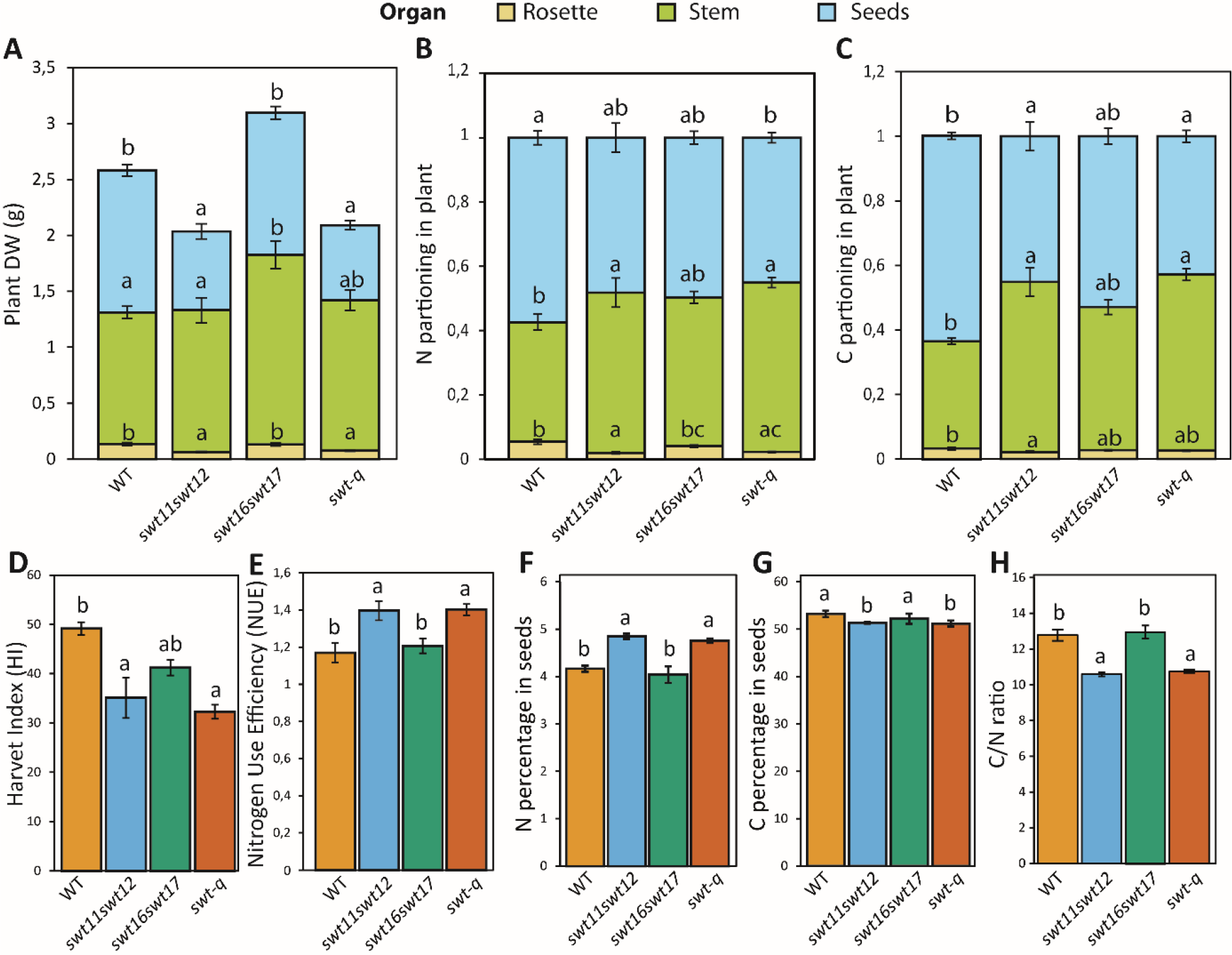
*SWEET* genes modulate plant biomass, yield and nitrogen use efficiency. Barplots showing the dry weight (A) and the C/N ratio (B) in different plant parts (i.e. rosette leaves, total stem and seeds) of wild type, *swt11swt12*, *swt16swt17* and quadruple mutants. Barplots showing the plant nitrogen use efficiency (NUE) (C) and the harvest index (HI) (D) for each genotype. Means +/− SE are shown (n≥5 for each genotype). A one-way ANOVA combined with the Tukey’s comparison post-test have been made to compare the different genotypes. The different letters indicate significant difference.

We also estimated the harvest index (HI) which corresponds to the ratio of seeds DW to aboveground plant DW at harvest time. In our growth conditions, HI represents about 50% in wild-type plants (Fig. 3D). It was significantly lower (by about 70%) in the *swt11swt12* and *swt-q* mutants while no significant difference was observed for the *swt16swt17* mutant compared to the wild type albeit a similar tendency (Fig. 3D). Since the *swt11swt12* and *swt-q* mutants produce less seeds (lower HI) with a lower proportion of N in seeds (lower NHI), we determined the Nitrogen Use Efficiency (NUE) as the ratio of Nitrogen Harvest Index (NHI)- to-Harvest Index (HI) to have information about the efficiency of the different lines to store N in seeds irrespective of the plant capacity to produced seeds. Interestingly, we measured an improved NUE in the *swt11swt12* and *swt-q* mutants compared to the wild type and the *swt16swt17* mutant (Fig. 3E). This suggest that *SWEET11* and *SWEET12* negatively impact the way the plant is efficiently using its nitrogen. Finally, we estimated the seed composition in the different genotypes and observed a significant decrease in the C percentage while the N percentage was significantly increased in the *swt11swt12* and quadruple mutants (Fig. 3F-G). As a consequence, the C:N ratio of the seeds was significantly decreased in both genotypes (Fig. 3H).

### Mutation in SWEET genes affects N remobilisation efficiency

Despite an improved NUE in the *swt11swt12* mutant, we observed a decreased C and N allocation to seeds in this genotype (Fig. 3B, C and E). In order to understand in which organ, the N flux might be blocked, we analysed the ^15^N partitioning to rosette, stem and seeds in the different mutants (Fig. 4). The use of ^15^N labeling allows to estimate the plant capacity to remobilize to the seeds the N compounds produced and stored in the rosette during the labeling period (Marmagne *et al.*, 2020). In wild-type plants, 5% of ^15^N remained in the rosette while 45% was stored in the stem, and 50% was remobilized to the seeds (Fig. 4A-C). Our results show that, consistently with a better NUE, less ^15^N remains in the rosette of the *swt11swt12* mutant (Fig. 4A). A similar tendency, albeit not significant, was observed in *swt16wt17* and quadruple mutants compared to wild-type plants (Fig. 4A). Interestingly, a significant increased proportion of ^15^N was measured in the stem of the *swt11swt12* and *swt-q* mutants (Fig. 4B) which led to a lower ^15^N remobilisation to the seeds (Fig. 4C). Finally, the calculation of the NRE index allowed to determine if the ^15^N partitioning to seeds is totally controlled by sink strength. In both the *swt11swt12* and *swt-q* mutants, we observed a significant increase of the NRE index compared to the wild-type plants (Fig. 4D). Altogether these results show that disruption of *SWEET11* and *SWEET12* gene expression leads to better nitrogen use efficiency and remobilisation efficiency from the rosette. However, the N remobilized from the rosette accumulates in the stem at the expense of the seeds, most probably explaining the lower harvest index.

**Fig. 4.**
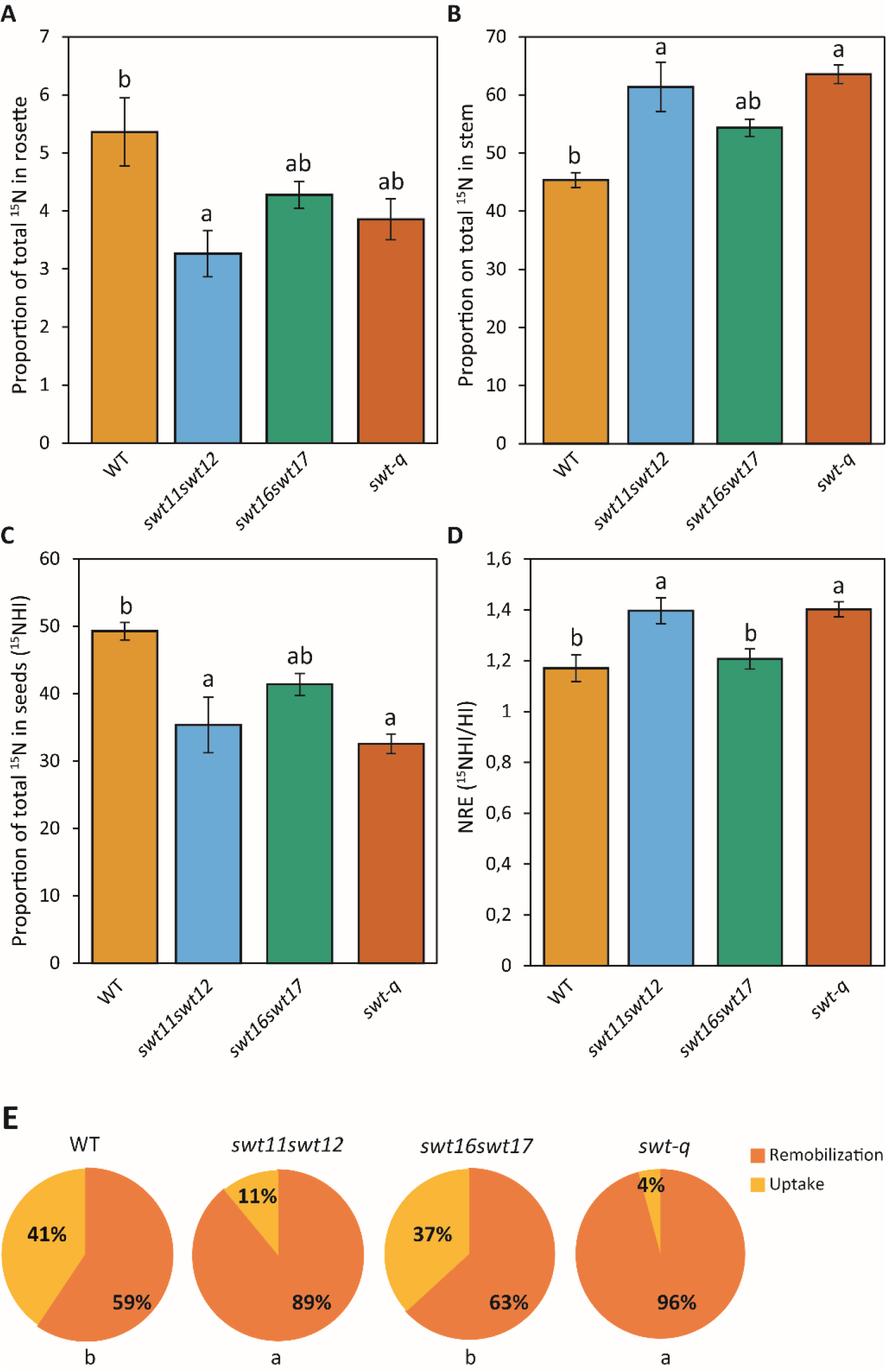
Nitrogen remobilisation efficiency is partly dependent on *SWEET* genes. Barplots showing the proportion on total ^15^N in rosette (A), stem (B), and seeds (C) and the N remobilisation efficiency (NRE) (D) in the wild-type, *swt11swt12*, *swt16swt17* and quadruple mutants. (E) Pie charts showing the proportion of N originating from N remobilisation or post-flowering N uptake in seeds of the wild type, *swt11swt12*, *swt16swt17* and *swt-q* mutants. Means +/− SE are shown (n≥5 for each genotype). A one-way ANOVA combined with the Tukey’s comparison post-test have been made to compare the different genotypes. The different letters indicate significant difference.

The nitrogen presents in seeds is coming from both N remobilisation and post-flowering N uptake by the roots. Based on the results of the ^15^N labeling, we could also deduce the proportion of both N origin in seeds (Marmagne *et al.*, 2022). In wild-type plants, about 60% of N found in seeds is coming from the remobilisation process while the remaining 40% is linked to the post-flowering N uptake (Fig. 4E). Interestingly, in the *swt11swt12* double mutant and in the *swt-q* mutant, these proportions are severely impacted and the N part originated from post-flowering uptake is strongly reduced (Fig. 4E). In the *swt16swt17* double mutant seeds, the N coming from the post-flowering uptake is also slightly modified compared to wild-type seeds but the difference is not statistically different (Fig. 4E). These results further strengthen the existence of an association of sugar transport with N uptake and remobilisation to seeds.

### Quantitative analysis of the different vascular bundle types and impact of mutations in SWEET genes

Nitrate, and amino acids are mostly transported in the xylem sap from leaves to seeds through the stem (Tegeder and Masclaux-Daubresse, 2018). Since we observed fewer xylem tissues in the stem of mutant lines together with an improved nitrogen remobilisation, we further explore the stem xylem defect by performing a quantitative analysis of the xylem in a vascular-bundle type manner (Fig. 5, and Supplementary Figs. S2-S5). Indeed, three different VB types can be observed in Arabidopsis stem: the M (main stem)-type VBs which are the most common and are characterised by a triangular shape; the B (branch)-type VBs that are dividing VBs connected to a lateral inflorescence stem; and the L (leaf)-type VBs that are round-shaped VBs connected to the cauline leaf vasculature (Fig. 5A). The M-type and B-type correspond to VBs involved in root-to-shoot transport of water and N derived nutrients to the seeds (Park *et al.*, 2015). In wild-type plants, we observed around 10 VBs among which between 4-6 M-type VBs, 2-3 B-type VBs and 2-3 L-type VBs (Fig. 5B). When the expression of *SWEET11* and *SWEET12* is disrupted, no significant change in the total number of VBs is observed (Fig. 5B). However, when the expression of both genes coding for the tonoplastic SWEET16 and SWEET17 transporters is impaired, a significant decrease of the total number of VBs is observed compared to the wild-type plants (Fig. 5B). Interestingly, in the *swt-q* mutant line, similarly to the *swt16swt17* mutant, we also found fewer VBs compared to the wild-type plants (Fig. 5B). While the same number of B-type VB is observed in the *swt16swt17* double mutant compared to the wild type, significantly fewer M-type VBs and a tendency for more L-type VBs are observed in this mutant line (Fig. 5B). In the *swt-q* mutant a trend, albeit not significant, for less M-type and L-type VBs and more B-type VBs is observed (Fig. 5B).

**Fig. 5.**
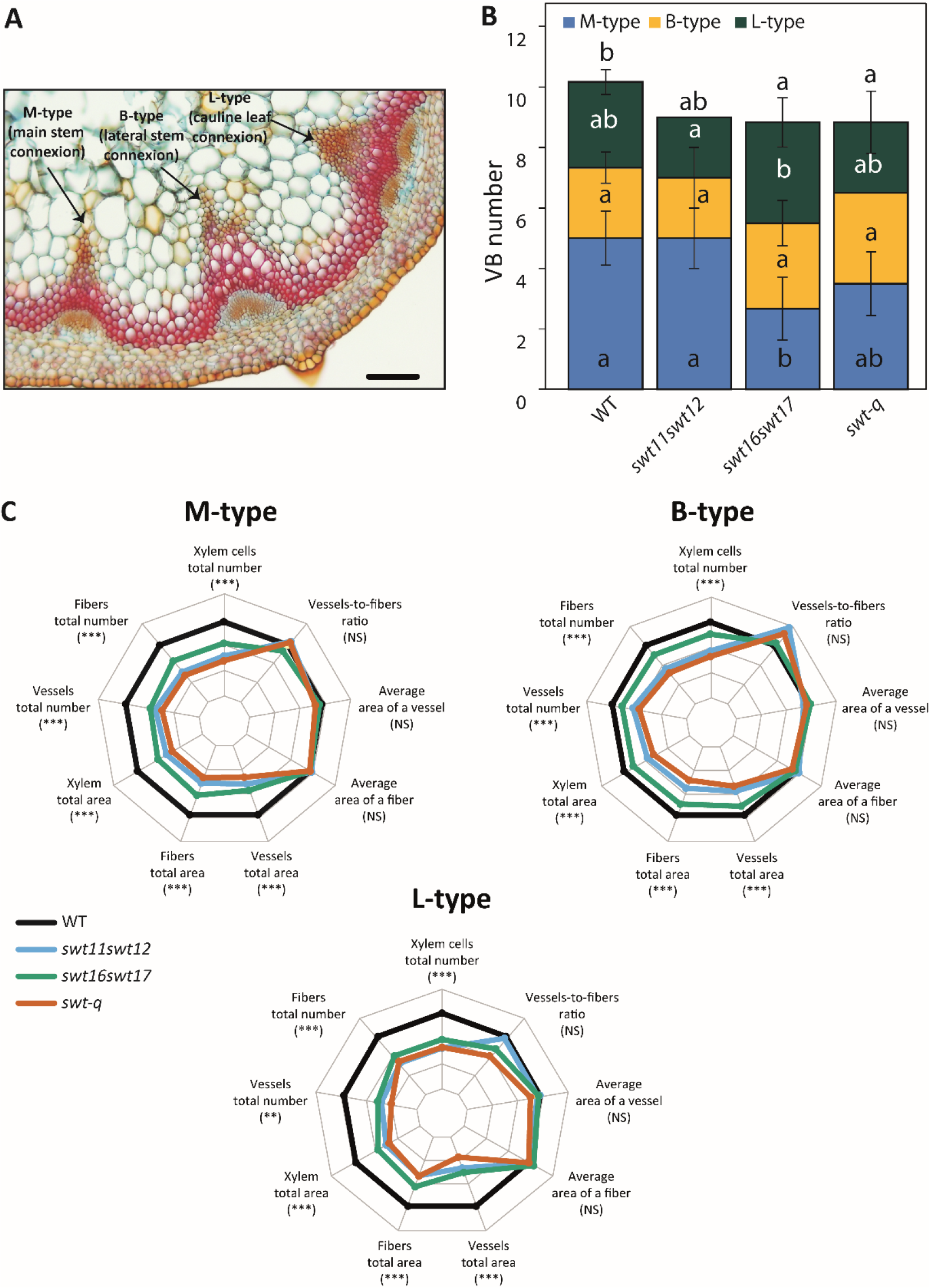
Mutation in *SWEET* genes affects the vascular bundle development. (A) Photograph of the different vascular bundle (VB) types within an inflorescence stem cross-section showing the M-type, B-type and L-type VBs. Scale bar = 100 μm. (B) Stacked histogram showing the number of each VB type in the different genotypes. Values are means ± SD (n=5-6 plants). (C) Heatmap of anatomical traits describing the xylem tissue in M-type, B-type and L-type VBs of the wild type, *swt11swt12*, *swt16swt17*, and *swt-q* mutants. The data are presented as the value of mutant lines over the value of WT plants ratios in different degree of blues for the statistically significant differences. The non-significant differences are presented in white. A one-way ANOVA combined with the Tukey’s comparison post-test has been performed on the raw data to compare the genotypes. The results of the ANOVA test are presented beside each VB type and those of the Tukey’s comparison post-test are presented in Supplementary Figs. 3, 4 and 5.

Several anatomical parameters such as xylary fibres and xylem vessels numbers and sizes were then measured to detail the morphological features of the different VB types. A principal component analysis (PCA) was applied to the dataset obtained on wild-type plants and allowed to observe a separation between the different VB types within the projection of the two first principal component planes that gather more than 83% of the total variation (Supplementary Fig. S2A). The first dimension clearly separates the B-type VB from the L-type VB while the M-type VB present an intermediate phenotype (Supplementary Fig. S2A). The number of xylem cells (xylary fibres and xylem vessels) and the area occupied by xylem vessels are the factors which are contributing the most to this separation (Supplementary Fig. S2B). The data show that B-type VBs are characterised by more and bigger xylary fibres than M-type VBs (Supplementary Fig. S2B). L-type VBs display fewer and smaller xylem vessels as well as a smaller fibre-to-vessel ratio than the M-type VBs (Supplementary Fig. S2B). In the mutant line, independently of the VB type, the *swt11swt12* and *swt-q* mutant lines displayed significantly fewer and smaller xylary fibres and xylem vessels (Fig. 5C and Supplementary Figs. 3, 4 and 5) compared to the wild type. Nonetheless, one exception is observed in the B-type VBs of the *swt11swt12* double mutant for which no difference of the total number of vessels is observed compared to the wild type (Supplementary Fig. 3C). In addition, we observe that, compared to the wild type, *swt16swt17* double mutant displayed significantly fewer and smaller xylem cells (including fibres and vessels) in M- and L-type VBs, but no significant effect was measured for B-type VBs (Fig. 5C and Supplementary Figs. 3, 4 and 5).

### SWEET11, SWEET12, SWEET16 *and* SWEET17 *have partially overlapping expression pattern in the vascular tissues of the stem independently of the VB type*

Next the translational GUS fusions lines previously described (Chen *et al.*, 2012; Guo *et al.*, 2014) were used to check the expression pattern of the different SWEET transporters in the different VB types (Fig. 6). We observed that all four transporters are expressed both in phloem and xylem whatever the VB type (Fig. 6). In the xylem tissue, SWEET11, SWEET16 and SWEET17 are expressed in the developing xylem cells (Figure 6B, H and K) while an expression of SWEET11, SWEET12 and SWEET17 is detected in the xylem parenchyma cells located at the bottom of the vascular bundle in the protoxylem area (Fig. 6A-C, D-F and J-L). This expression pattern is particularly visible in L-type VBs which are mainly composed of parenchyma cells (Fig. 6C, F and L). In addition, SWEET11 and SWEET12 are localised in the starch sheath cells, situated between the cortex and the phloem (Altamura *et al.*, 2001) (Fig. 6A-F). An expression of SWEET16 and SWEET17 is observed in the cortex cells and the interfascicular fibers (Fig. 6G-L). Finally, SWEET17 is also expressed in some of the pith cells (Fig. 6J-L).

**Fig. 6.**
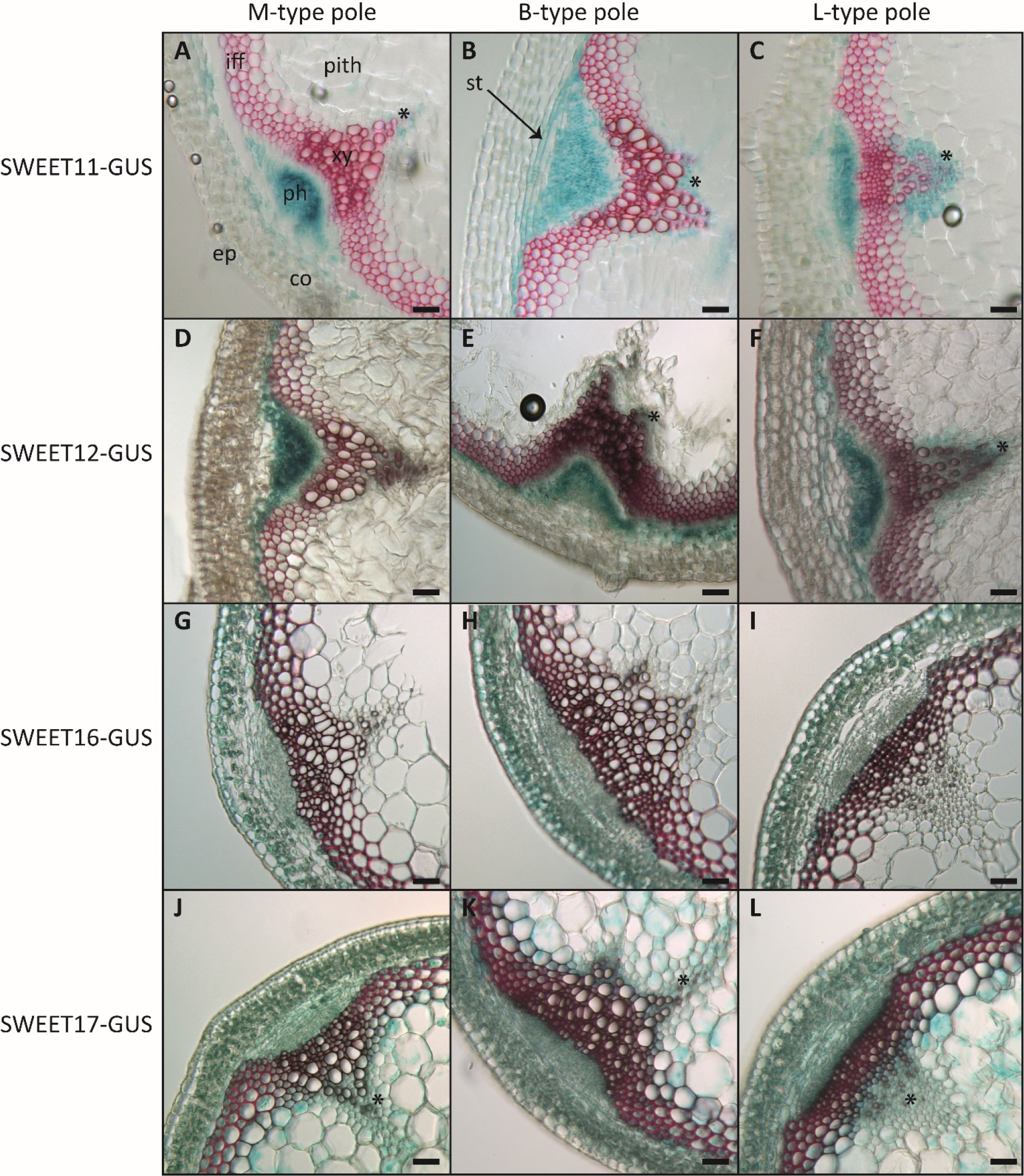
Expression pattern of SWEET11, 12, 16 and 17 in the different vascular bundle types and adjacent tissues. Histochemical analysis of GUS activity in lines expressing SWEET11-GUS (A to C), SWEET12-GUS (D-F), SWEET16-GUS (G-I) or SWEET17-GUS (J-L) fusion proteins driven by their native promoter in sections taken at the bottom of the inflorescence stem after 7-weeks of growth. Photographs are showing GUS expression in M-type vascular bundle (A, D, G and J), in B-type vascular bundle (B, E, H and K) or in L-type vascular bundle (C, F, I and L). Asterisks point to cells showing blue GUS staining in xylem parenchyma cells. Lignin is colored pink after phloroglucinol staining. ep, epidermis; co, cortex; iff, interfascicular fibers; ph, phloem; st: starch sheath cells; xy, xylem.

## Discussion

Modulation of inter or intracellular pool of sugars by modifying the expression of genes involved in sugar metabolism or transport modifies shoot and root growth, and plant yield in several species (e.g., Arabidopsis, Tomato, Tobacco, Poplar and Pea) (Dai *et al.*, 1999; Park *et al.*, 2008; Wingenter *et al.*, 2010; Zhang *et al.*, 2010; Mahboubi *et al.*, 2013; Le Hir *et al.*, 2015; Stein *et al.*, 2017; Lu *et al.*, 2020; Valifard *et al.*, 2021; Aubry *et al.*, 2022). However, the effect of simultaneous disruption of sugar transport at both inter and intracellular levels has not been explored so far. In this study we explored the effect on plant development and biomass production of the disruption of two genes coding for proteins involved in intercellular sugar transport, SWEET11, SWEET12, together with two genes coding for intracellular sugar transporters, SWEET16 and SWEET17. *SWEET11* and *SWEET12* genes encode sugar transporters located at the plasma membrane of vascular parenchyma cells in leaves and stem (Chen *et al.*, 2012; Le Hir *et al.*, 2015; Cayla *et al.*, 2019; Kim *et al.*, 2021b). They were also shown to be expressed in seeds and roots (Chen *et al.*, 2015; Desrut *et al.*, 2020). On the other hand, *SWEET16* and *SWEET17* encode tonoplast-localized sugar transporters expressed in the vascular system of leaves, stem and roots (Chardon *et al.*, 2013; Klemens *et al.*, 2013; Guo *et al.*, 2014; Valifard *et al.*, 2021; Aubry *et al.*, 2022). Comparing the expression pattern of the four SWEET transporters in inflorescence stem, we observed that they have partially overlapping expression pattern in the phloem, the developing xylem cells and the xylem parenchyma cells. In addition, SWEET11 and SWEET12 are expressed in endodermis-like cells which transiently store starch (also called starch sheath cells) (Altamura *et al.*, 2001) and SWEET16 and SWEET17 in the cortex cells. Interestingly the phenotype of the quadruple mutant, for most of the traits related to plant biomass production and plant yield, is similar to that of the *swt11swt12* double mutant. On the other hand, the phenotype of the *swt16sw17* double mutant was similar to that of the wild type or intermediate between the *swt11swt12* and *swt-q* mutants. These results suggest that intercellular facilitated transport of sugars, mediated by SWEET11 and SWEET12, expressed conjointly in the starch sheath cells, the phloem and, the xylem parenchyma, constitutes one limiting factor for plant biomass production and plant yield. On the other hand, the maintenance of the sugar homeostasis within the cell, regulated by SWEET16 and SWEET17 constitutes an important parameter for the regulation of the xylem and interfascicular fibres development, consistently with their expression in these cell types (Aubry *et al.*, 2022).

In Arabidopsis, the inflorescence stem results from two distinct morphogenetic transitions, the so-called bolting transition and floral transition (Pouteau and Albertini, 2009). The first part of the stem (cauline leaf zone), resulting from the bolting transition, is composed of cauline leaves carrying an axillary flowering shoot while the “true inflorescence” carries the flowers and siliques. Interestingly, we show that the reduction of inflorescence stem height in the *swt11swt12* double mutant is mainly due to a specific decrease of the “true inflorescence” height, suggesting that both transporters play a role in the floral transition. It is known that during the floral transition, an increase of the sucrose in the phloem sap is required (Corbesier *et al.*, 1998) and that the primary metabolism is reprogrammed including the transcriptional activation of the gene coding for the sucrose transporter *SWEET10* by FLOWERING LOCUS T (FT) transcription factor (Andrés *et al.*, 2020). Although a similar regulation of *SWEET11* and *SWEET12* expression by FT is unlikely according to Andrés et al. (2020), the defect in the sucrose phloem loading, previously observed in the source leaves of the *swt11swt12* double mutant (Chen *et al.*, 2012) could account for this phenotype.

In the stem, we also show a perturbation of the C partitioning (including an accumulation of soluble sugars and starch) along with an expression of both SWEET11 and SWEET12 in the stem phloem tissue. These suggest that both transporters play a role in loading the phloem in the stem as previously shown in leaves (Chen *et al.*, 2012). Moreover, our results show that in stem they are also likely involved in the starch remobilisation from the starch sheath cells to the phloem parenchyma cells. These results are in line with stems being a major contributor to lifetime carbon gain during plant development (Earley *et al.*, 2009). Especially, it has been shown that before the first siliques development, the main inflorescence stem store carbon such as starch mainly in the endodermis-like cell layer above the phloem cells. Later in their development, stems are re-exporting nutrients to sustain siliques development and seeds maturation (Altamura *et al.*, 2001; Ohmae *et al.*, 2013; Sugita *et al.*, 2016; Durand *et al.*, 2018). However, to which extent perturbations of the C allocation in stems impacts C percentage in seeds is unclear. Part of the answer is provided by the C and N percentages in stems and seeds obtained in Chardon et al. (2014) on an Arabidopsis RIL population. Interestingly, while seed N% is positively correlated with stem N%, there is no significant correlation between seed C% and stem C%. These results suggest that C remobilisation from the stem is much less important for seed filling than the N remobilisation. Nonetheless, our results support that the lower C% in seeds measured in the *swt11swt12* mutant could partially result from the overall defective sugar loading in this mutant. This will lead to a decrease delivery of carbon to the seeds because part of the C is not properly loaded or remobilized in the phloem sap from the stem and accumulated as starch in this organ. In addition, both *SWEET11* and *SWEET12* together with *SWEET15* have also been shown to be expressed in the different tissues of the seed (i.e., micropylar, endosperm and seed coat) across its development (Chen *et al.*, 2015). Consistently *swt11swt12* and *swt11swt12swt15* mutant seeds display a decrease in their total fatty acid content (Chen *et al.*, 2015). Therefore, the lower C% in the double mutant seeds could also be the consequence of the defect in sugar transport within the different tissues of the seed. In the future, a seed-specific complementation of the mutant phenotype could help to distinguish between the contribution of the stem or seed in seed carbon allocation defect.

In agriculture, one of the dilemmas to solve in the near future consists in improving the plant nitrogen use efficiency (NUE) to reduce production costs and environmental risks linked to N leakage in the environment. While manipulating the expression of amino acid transporters has been shown to efficiently improve NUE (Yang *et al.*, 2020), the effect of a deregulation of sugar transporters on NUE is scarce (Schofield *et al.*, 2009; Klemens *et al.*, 2013). This is surprising since C availability is known to be strongly associated with NUE (Fernie *et al.*, 2020). In this study, we show that mutation in the expression of both *SWEET11* and *SWEET12* impact the N allocation at the whole plant level. Indeed, we measured (i) an increase of amino acids content in the stem, (ii) an increase of N partitioning to the stem concomitantly with a decrease in the seeds and in the rosette leaves, and (iii) an increase of ^15^N proportion in the stem and decrease in the seeds. Ultimately, *swt11swt12* double mutant displays an improved NUE and NRE despite a lower harvest index. Finally, *swt11swt12* seeds display a lower C:N ratio which points to a non-proportional enhancement of nitrogen supply (higher N%). In seeds of several species such as sunflower, rapeseed, soybean and Arabidopsis, both C and N percentages are strongly correlated (Marmagne *et al.*, 2020). Up to now, three hypotheses have been proposed to explain this phenomenon: (i) the dilution effect of the protein concentration by the enhanced oil concentration and *vice versa*, (ii) the competition effect between C and N metabolisms for energy, and (iii) the N availability effect which suggests that the local N quantity and quality may affect the C and N metabolisms in siliques (Marmagne *et al.*, 2020). In addition, we could also propose that the retention of C and N in *swt11swt12* stem could also be responsible for this lower C:N ratio in seeds. In this context, a feedback mechanism consisting in a better nitrogen remobilisation efficiency from the rosette leaves is used to compensate for the partial retention of N in stem. Moreover, our results point out that the *swt11swt12* double mutant also displays a defect in the post-flowering N uptake by the roots. Interestingly, in Arabidopsis seedlings, Chen et al. (2016) identified that the regulation of the expression of *SWEET11* and *SWEET12* by the shoot-to-root mobile transcription factor ELONGATED HYPOTOCOL5 (HY5) is required to modulate the expression of the nitrate transporter *NRT1.2* in roots. Our results support this model and further suggest that SWEET16 and SWEET17 could also be considered since the *swt-q* mutant displays a tendency towards a further decrease in post-flowering N uptake albeit not significant. Consistently, *swt17* single mutant showed defects in root growth and architecture (Valifard *et al.*, 2021) that could also account for N uptake impairment since both traits are strongly linked (Kiba and Krapp, 2016). Altogether our results suggest that a proper sugar phloem loading mediated by SWEET11 and SWEET12 negatively impacts the way the plant is using and remobilising its nitrogen to the seeds. However, to better apprehend the C and N fluxes in the different *swt* mutant lines, ^13^C and ^15^N labelling studies together with phloem and xylem saps composition measurements could be of interest.

To be transported at long-distance, nutrients largely rely on the plant vascular system. Especially, the inflorescence stem constitutes the obligatory way for nutrients to be distributed from rosette leaves to the seeds (van Bel, 2021). Consistently with our previous works on *swt11swt12* and *swt16swt17* double mutant lines (Le Hir *et al.*, 2015; Aubry *et al.*, 2022), we further show that the disruption of the expression of the four *SWEET* genes affects the vascular system expansion and tissue distribution. Especially the stem diameter decrease is likely due to the impaired development of cells undergoing important secondary cell wall formation (i.e., xylem and interfascicular fibers). As described in Park et al. (2015), three different types of VB with distinct functional features regarding water and mineral transport can be found in Arabidopsis stem. In this study, we show that the VB types also display distinct anatomical features, consisting in different proportion of xylary fibres, xylem vessels and xylem parenchyma cells. These observations suggest the existence of a genetic control of their differentiation, which will most probably involved transcription factors responsible for the xylem cell type differentiation together with hormone regulation (Schuetz *et al.*, 2012). Nonetheless, since sugars have also been shown to affect xylem development (Aloni, 1987), we hypothesize that changes in the sugar availability mediated by SWEET transporters, could be at play in the VB type differentiation. In line with this hypothesis, a lower number of VBs, mainly due to less M-type VBs, was found only in *swt16swt17* mutant, along with a modified cellular morphology in the VBs involved in the connexion with the cauline leaves (L-type VB) and those responsible for long-distance transport of water and nutrients (M-type VB) (Park *et al.*, 2015). Overall, these results suggest a possible reduced capacity for water and nutrient transport as well as reduced exchanges of these compounds between the stem and the cauline leaves that should be further examined. Nonetheless, since the plant yield is not affected in this mutant, it is likely that genetic compensation occurs to sustain normal seeds quantity and quality. Therefore, the intracellular sugar fluxes mediated by SWEET16 and SWEET17 specifically affect the stem xylem development without impacting the plant yield in normal growth conditions. It could be of interest to test if the same holds true when plants are grown in limiting conditions such as osmotic stress for which xylem transport is important (Shafi *et al.*, 2015; Shinohara *et al.*, 2019). On the other hand, the defect in the sugar export/remobilisation capacity in the stem likely account for the global stem growth and development phenotype observed in the *swt11swt12* double mutant. Then, the resulting increased sugar storage in stem and leaves disturbs the plant capacity to deal with N and *in fine* the plant yield.

Interestingly, a significant negative correlation was found between stem diameter and nutrient contents (i.e. soluble sugars, starch and amino acids). Considering that stem diameter and nutrient contents could account for xylem development and C/N allocation respectively, our results support a link between both traits. A role for sugars in the vasculature development has been proposed for long, even if the identification of molecular actors involved starts only to emerge (for review Dinant and Le Hir, 2022). On the other hand, a similar role for amino acids has not been explored so far. Nonetheless, in tree stem, a nitrogen addition, an increased carbon allocation or, to a lesser extent phosphorus addition, promotes the enlargement of the vascular system (Plavcová *et al.*, 2013; Cai *et al.*, 2017; Hacke *et al.*, 2017; Hartmann *et al.*, 2020). At the opposite, in sunflower root, a high root N has been associated with lower vessels number and size in sunflower (Bowsher *et al.*, 2016). Thus, this organ-dependent relationship between vascular development and C/N allocation highlights the complex interaction between both traits. A higher complexity level must also be added since nutrients act both as energy-provider compounds and signal molecules (Fichtner *et al.*, 2021). Therefore, future studies exploring the link between nutrients allocation and vascular system development must consider a multiscale approach from tissue to whole plant level.

## Supplementary data

The following supplementary data are available at JXB online.

*Fig. S1.* Molecular characterisation of the quadruple *sweet* mutant lines.

*Fig. S2.* Distinct morphological features of the different vascular bundle’s type in wild-type stems.

*Fig. S3.* Cellular morphology of M-type pole in the different genotypes. *Fig. S4.* Cellular morphology of B-type pole in the different genotypes.

*Fig. S5.* Cellular morphology of L-type pole in the different genotypes.

*Table S1.* Primers used for characterising the quadruple mutant line.

## Acknowledgments

We thank Catherine Bellini for critical reading of the manuscript.

## Authors contribution

Conceptualization, R.L.H.; investigation, B.H., E.A., A.M., F.C. and R.L.H.; methodology, B.H., E.A. and A.M.; visualization, S.D., F.C. and R.L.H.; writing—original draft, R.L.H.; writing—review and editing, S.D., F.C. and R.L.H.

## Conflicts of interest

The authors declare that the research was conducted in the absence of any commercial or financial relationships that could be construed as a potential conflict of interest.

## Funding

This work has benefited from the support of IJPB’s Plant Observatory technological platforms and from a French State grant (Saclay Plant Sciences, reference ANR-17-EUR-0007, EUR SPS-GSR) managed by the French National Research Agency under an Investments for the Future program (reference ANR-11-IDEX-0003-03) through PhD funding to E.A.

## Data availability

The data supporting the findings of this study are available from the corresponding author, Rozenn Le Hir, upon request.

## Notes

### Competing Interest Statement

The authors have declared no competing interest.

### Summary of Updates

Supplementary data added to the manuscript

