## Supplementary files for "Impairment of sugar transport in the vascular system acts on nitrogen remobilisation and nitrogen use efficiency in Arabidopsis"

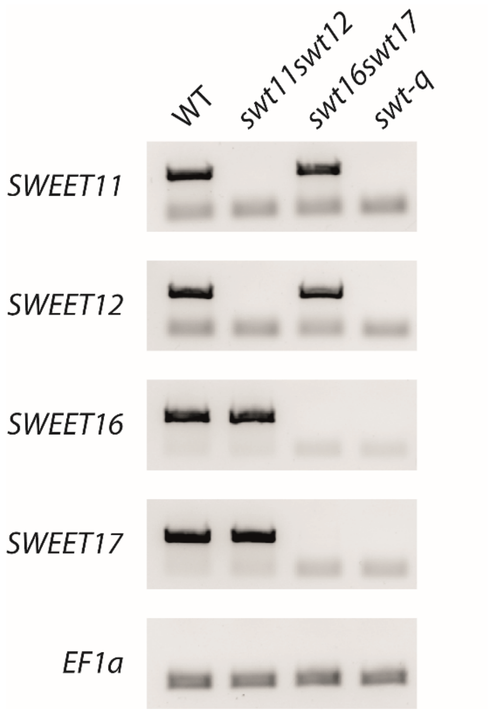

**Supplementary Fig. S1. Molecular characterisation of the quadruple *sweet* mutant lines.** RT-PCR analysis of *SWEET11*, *SWEET12*, *SWEET16* and *SWEET17* expression in the floral stem of the *swt11swt12*, *swt16swt17* and *swt-q* mutant lines. Total RNA was isolated from 7-week-old floral stem of plants grown in long-day conditions. After reverse transcription, the cDNAs were used for amplification with primers designed between the start and stop codon of the *SWEET11*, *SWEET12*, *SWEET16* and/or *SWEET17* sequences (for primers sequences see Supplemental Table 1). Expression of *EF1α* was used as a loading control.

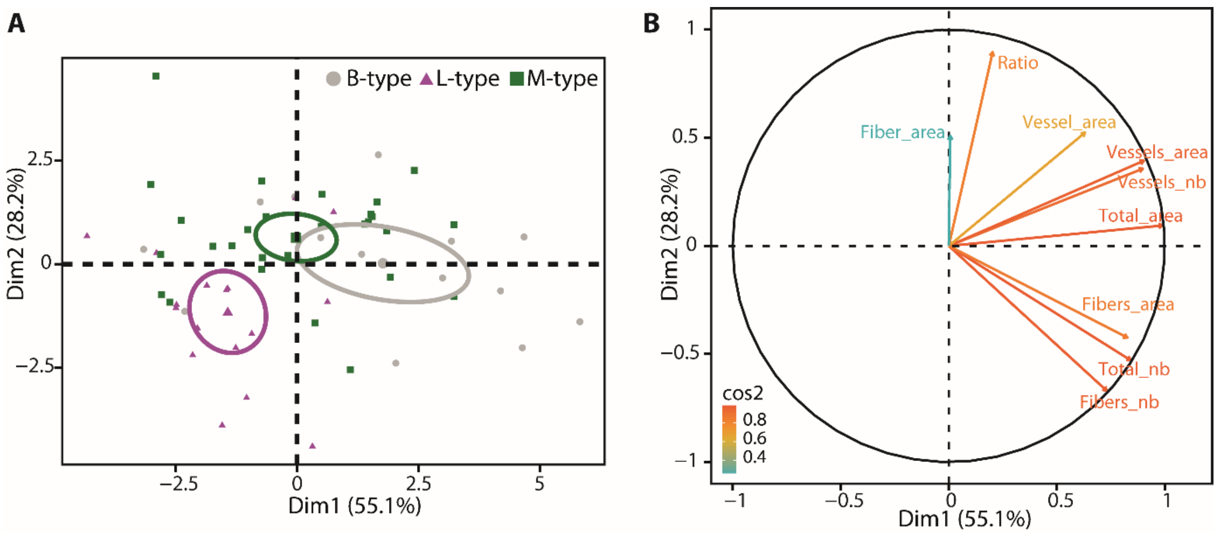

**Supplementary Fig. S2. Distinct morphological features of the different vascular bundle's type in wild-type stems.** (A and B) Principal component analysis of anatomical factors measured in the different VB type in wild-type stems. The PCA is showing the projection on the 2 first components of the samples (namely the different VB type) (A) and factors (B). The different factors are the number of xylem cells (Total_nb), of xylary fibers (Fibers_nb), of xylem vessels (Vessels_nb) and the ratio xylem vessels-to-xylary fibres number (Ratio) per VB type; the average area occupied by xylem cells (Total_area), by xylary fibers (Fibers_area) and by xylem vessels (Vessels_area) and the average size of a xylary fiber (Fiber_area) and of a xylem vessel (Vessel_area). In (B) the different factors are color-coded on a red-to-green scale depending on their contribution to the PCA as provided by the cos2 value. For raw data, see the Supplementary Dataset S1.

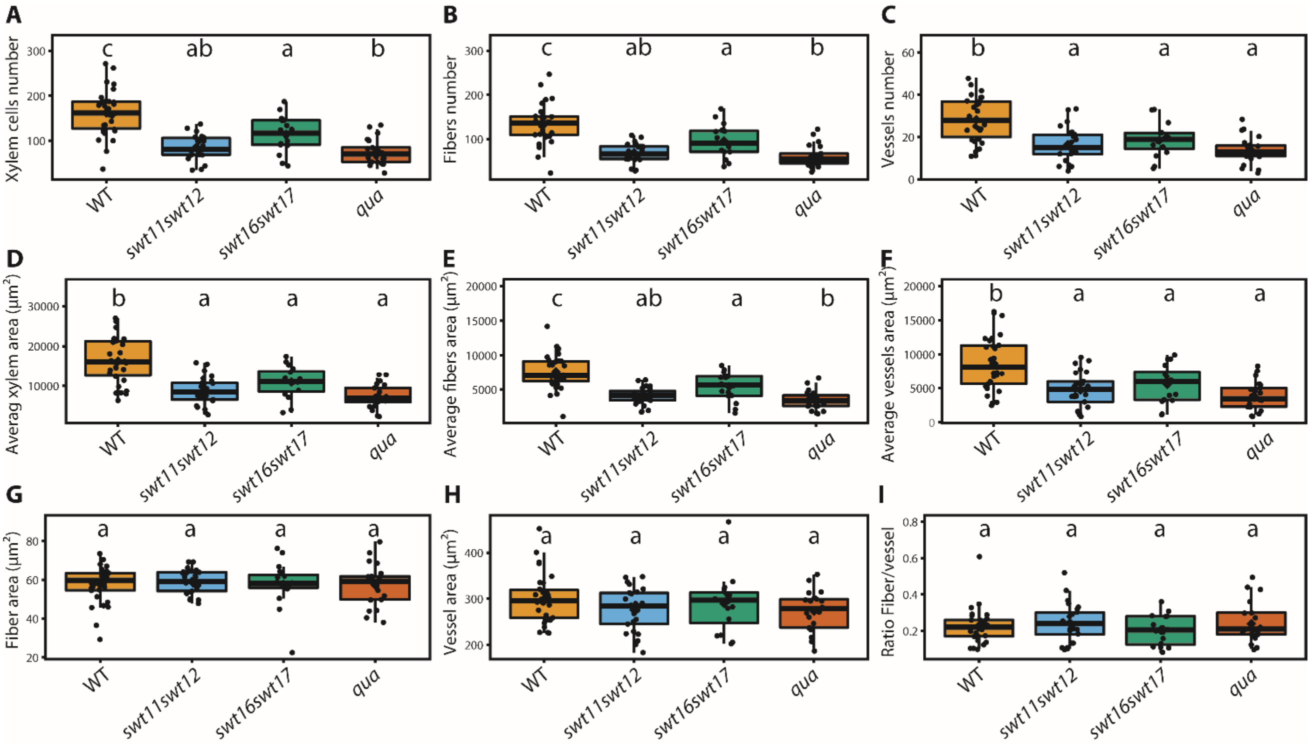

**Supplementary Fig. S3. Cellular morphology of M-type pole in the different genotypes.** Boxplots showing the total number of xylem cells (A), the average number of xylary fibers cells per VB (B) and the average number of xylem vessels (C) per VB type. Average area occupied by xylem tissue (D), average area occupied by xylary fibers (E) and average area occupied by xylem vessels (F). Average area of an individual xylary fiber (G) or an individual xylem vessel (H) as well as the fiber-to-vessel ratio (I). The box-and-whisker plots represents values from 30, 16, 25 and 21 M-type poles for wild type, *swt11swt12*, *swt16swt17* and quadruple mutant lines, respectively. The lines represent median values, the tops and bottoms of the boxes represent the first and third quartiles, respectively, and the ends of the whiskers represent maximum and minimum data points. A one-way ANOVA combined with the Tukey's comparison post-test have been made to compare the pole types. The different letters indicate significant difference ($P < 0.05$).

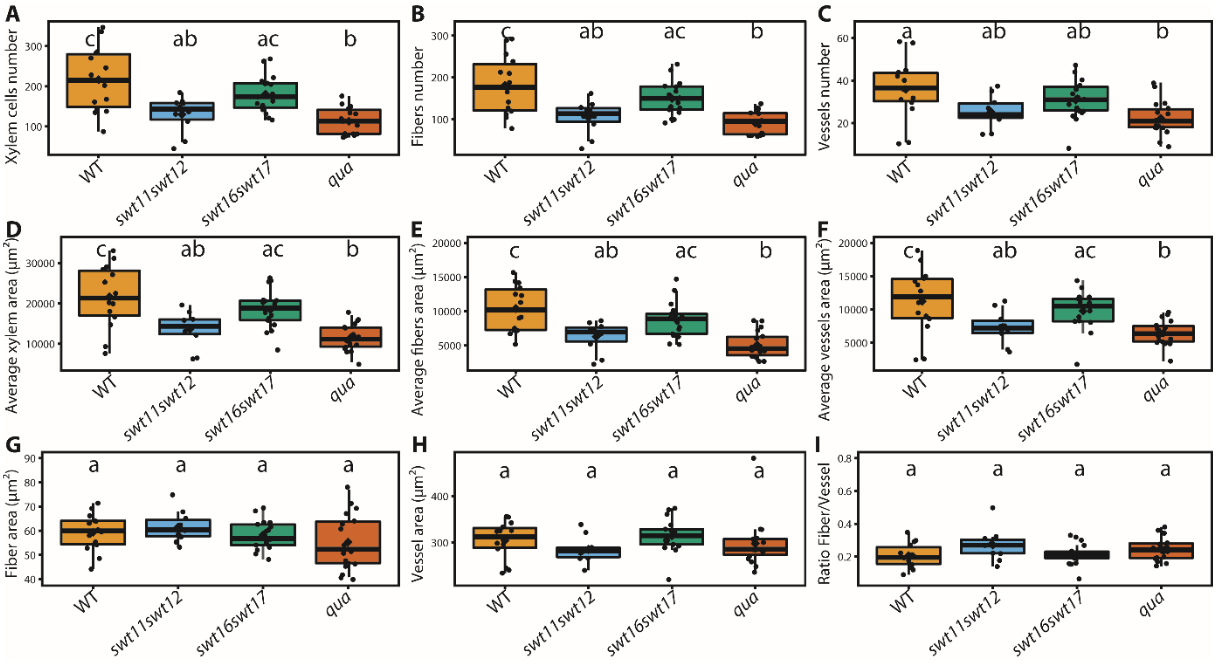

**Supplementary Fig. S4. Cellular morphology of B-type pole in the different genotypes.** Boxplots showing the total number of xylem cells (A), the average number of xylary fibers cells per VB (B) and the average number of xylem vessels (C) per VB type. Average area occupied by xylem tissue (D), average area occupied by xylary fibers (E) and average area occupied by xylem vessels (F). Average area of an individual xylary fiber (G) or an individual xylem vessel (H) as well as the fiber-to-vessel ratio (I). The box-and-whisker plots represents values from 14, 17, 10 and 18 B-type poles for wild type, *swt11swt12*, *swt16swt17* and quadruple mutant lines, respectively. The lines represent median values, the tops and bottoms of the boxes represent the first and third quartiles, respectively, and the ends of the whiskers represent maximum and minimum data points. A one-way ANOVA combined with the Tukey's comparison post-test have been made to compare the pole types. The different letters indicate significant difference ($P < 0.05$).

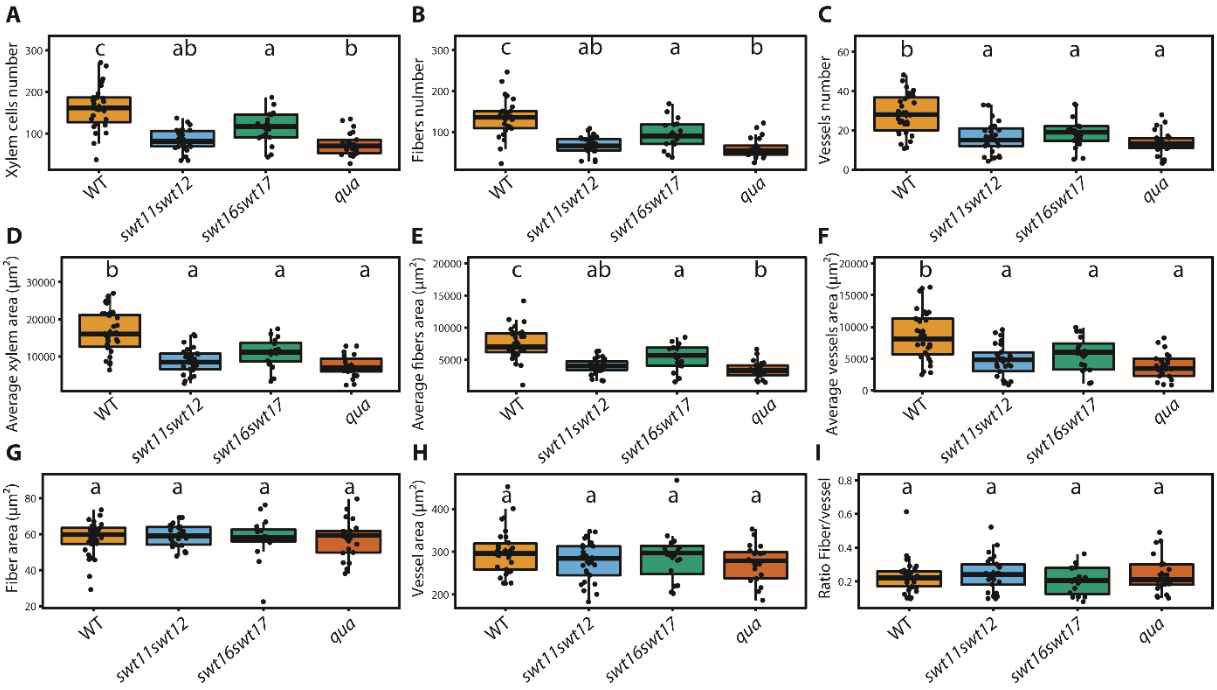

**Supplementary Fig. S5. Cellular morphology of L-type pole in the different genotypes.** Boxplots showing the total number of xylem cells (A), the average number of xylary fibers cells per VB (B) and the average number of xylem vessels (C) per VB type. Average area occupied by xylem tissue (D), average area occupied by xylary fibers (E) and average area occupied by xylem vessels (F). Average area of an individual xylary fiber (G) or an individual xylem vessel (H) as well as the fiber-to-vessel ratio (I). The box-and-whisker plots represents values from 17, 20, 10 and 13 L-type poles for wild type, *swt11swt12*, *swt16swt17* and quadruple mutant lines, respectively. The lines represent median values, the tops and bottoms of the boxes represent the first and third quartiles, respectively, and the ends of the whiskers represent maximum and minimum data points. A one-way ANOVA combined with the Tukey's comparison post-test have been made to compare the pole types. The different letters indicate significant difference ($P < 0.05$).

**Supplementary Table S1. Primers used for characterising the quadruple mutant line.**

| Accession number | Primer name | Sequence (5'→3') | Amplicon size (bp) | Purpose |
|---|---|---|---|---|
| At3g48740 | sweet11-1_LP | GATGTTTTGGAGACCAAGCTG | 1558 | Genotyping |
| | sweet11-1_RP | GGCTCCTTTATGGTCTTGCTC | | |
| At5g23660 | sweet12-1_LP | CTAAACCCGGAACCAAAGATC | 1638 | |
| | sweet12-1_RP | GAACAAACCCAATCACGTTTG | | |
| At3g16690 | sweet16-4_LP | TGCAAATAATTTAGCAACCGC | 1742 | |
| | sweet16-4_RP | TATAAATGATCTGGGGCCATC | | |
| At4g15920 | sweet17-1_LP | TGATGTGAGGCCTTCCTCTT | 771 | |
| | sweet17-1_RP | CCGTTTTGGTTGTCGTTTTT | | |
| At3g48740 | SWEET11ATG-stop_F | ATGAGTCTCTTCAACACTGAAAACAC | 870 | Full-length PCR |
| | SWEET11ATG-stop_R | TCATGTAGCTGCTGCGGA | | |
| At5g23660 | SWEET12ATG-stop_F | ATGGCTCTCTTCGACACTCATAA | 858 | |
| | SWEET12ATG-stop_R | TCAAGTAGTTGCAGCACTGTTTC | | |
| At3g16690 | SWEET16ATG-stop_F | ATGGCAGACTTGAGTTTTTATGTC | 693 | |
| | SWEET16ATG-stop_R | TTAAGCGAGGAGAGGTTGATTT | | |
| At4g15920 | SWEET17ATG-stop_F | ATGGCAGAGGCAAGTTTCTATATC | 726 | |

| | | |
|---|---|---|
| | SWEET17ATG-stop_R | TTAAGAGAGGAGAGGTTCAACACG |
